# Bidirectional valence coding in amygdala intercalated clusters: A neural substrate for the opponent-process theory of motivation

**DOI:** 10.1101/2024.06.18.599516

**Authors:** Kenta M. Hagihara, Andreas Lüthi

**Affiliations:** Friedrich Miescher Institute for Biomedical Research, Basel, Switzerland; University of Basel, Basel, Switzerland; Allen Institute for Neural Dynamics, Seattle, USA

## Abstract

Processing emotionally meaningful stimuli and eliciting appropriate valence-specific behavior in response is a critical brain function for survival. Thus, how positive and negative valence are represented in neural circuits and how corresponding neural substrates interact to cooperatively select appropriate behavioral output are fundamental questions. In previous work, we identified that two amygdala intercalated clusters show opposite response selectivity to fear- and anxiety-inducing stimuli – negative valence (Hagihara et al. 2021). Here, we further show that the two clusters also exhibit distinctly different representations of stimuli with positive valence, demonstrating a broader role of the amygdala intercalated system beyond fear and anxiety. Together with the mutually inhibitory connectivity between the two clusters, our findings suggest that they serve as an ideal neural substrate for the integrated processing of valence for the selection of behavioral output.

## Introduction

Certain objects and events within our environment have significant motivational importance due to their impact on our welfare, survival, and reproduction. Depending on the behavioral responses they trigger, these environmental stimuli can possess either positive (appetitive, rewarding) or negative (aversive, punishing) valence. In response to stimuli with positive valence, animals show approaching/hedonic behavior (Peterson et al. 1972); in response to negative valence, defensive behavior including avoidance/flight and freezing (Fanselow and Bolles 1979; R. J. Blanchard et al. 2008; D. C. Blanchard, Griebel, and Blanchard 2001; Tovote et al. 2016; Fadok et al. 2017). Animals are innately programmed to perceive certain stimuli as having a specific valence, but they can also learn the valence of originally neutral stimuli, often through associative learning processes (Pavlov 1927). Thus, to understand how stimuli with different valence, that can occur simultaneously, are represented and processed in neural circuits has been a fundamental research topic. Over the past decade, there has been considerable advancement in the identification of brain areas and circuits that underlie valence processing (see (Tye 2018) for review). However, the neuronal mechanisms that process and integrate opposing valences, and then select appropriate behavioral output selection as a result of a synergistic orchestration of distributed neural circuits (Gross and Canteras 2012; Tovote, Fadok, and Lüthi 2015; Herry and Johansen 2014), are not well understood.

A unique feature of valence processing is that positive and negative emotions are opposed to each other, as described by Solomon and Corbit (Solomon and Corbit 1974) in a theory called the “opponent-process theory of motivation”. According to this theory, the cessation or absence of an expected negative stimulus induces a positive affect, while conversely, the cessation or absence of an expected positive stimulus results in the opposite, negative affect. Although this theory is widely accepted in the field of psychology, the neural circuit-level mechanisms involved have not been well studied. In our previous study, we characterized the role of the amygdala intercalated clusters (ITCs, see (Asede, Doddapaneni, and Bolton 2022) for a recent review) in fear and anxiety and found that: 1) two major clusters, the dorsomedial ITC (ITCdm) and the ventromedial ITC (ITCvm) cluster, mutually inhibit each other via GABAergic monosynaptic connections; 2) ITCdm is strongly activated by aversive foot shocks, whereas ITCvm does not respond to foot shocks but rather increases its activity when expected foot shocks are omitted (Hagihara et al. 2021). Thus, the anatomical and functional opponency makes the ITC system an ideal neural substrate for implementing the opponent-process theory. To further test this idea, in the current study, we extended our in vivo analysis and examined value intensity and positive valence coding in the ITC system.

## Results

To assess the properties of the same ITC neurons in response to multiple stimuli with opposite valence, we chronically monitored the activity of individual ITC neurons in freely moving mice using *in vivo* deep brain calcium imaging with a miniaturized microscope (Ghosh et al. 2011; Hagihara et al. 2021) (**Fig. 1a-c**). First, we investigated whether the ITC_dm_ neurons represent the value of aversive stimuli, i.e., scale their responses as a function of shock intensity. Mice were presented five times with mildly aversive foot shock stimuli of five different intensities (**Fig. 1c,d**). We found that a large fraction of ITC_dm_ neurons (78.7%) were excited by foot shocks at 0.65 mA, which is commonly used for fear conditioning. Individual ITC_dm_ neurons showed reliable responses to shocks of the same intensity across trials in terms of response amplitude and latency from shock onset (**Fig. 1e-g**), suggesting reliable information transmission from the peripheral pain sensing system. Population-averaged ITC_dm_ responses increased with larger shock intensity (**Fig. 1f-I**, *P* = 7.7 × 10^−6^, Jonckheere-Terpstra test, n = 127 neurons), while the proportion of significantly responding neurons was consistent across shock intensities (**Fig. 1j**). This indicates individual neurons scale their activity to represent negative stimulus value, but that response probability remains constant at the population level. Moreover, even in a mouse from which we collected data over 3 days, response patterns were stable (**Fig. 1f**), suggesting that the upstream circuits that convey shock signals to ITC_dm_ are stable over time, as is the excitability of ITC_dm_ neurons.

**Figure 1.**
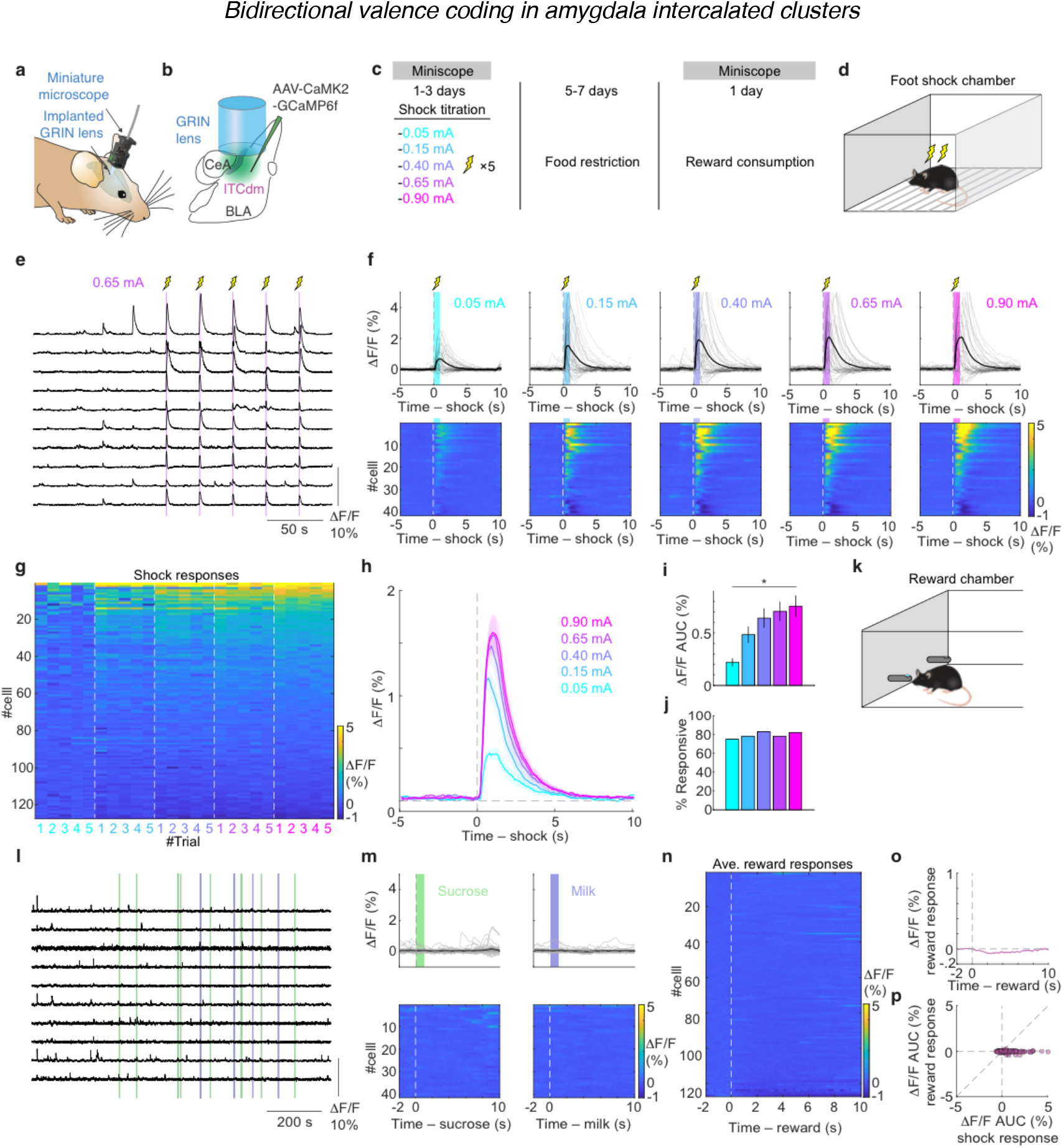
ITC_dm_ valence coding. **a**, Endoscopic imaging with a miniaturized microscope in a freely-moving mouse. **b**, AAV encoding CaMK2-GCaMP6f targeted to CeA, BLA and ITC_dm_. GRIN lens implanted above injection site. **c**, Experimental time course. **d**, Experimental scheme for shock titration experiment. **e**, Example Ca^2+^ traces. **f**, Trial-averaged ΔF/F Ca^2+^ time-course aligned to shock onset of all recorded ITC_dm_ from a representative mouse. Cells were sorted based on their shock response to 0.90mA. **g**, Area under the curve (AUC) ΔF/F values from all recorded ITC_dm_ neurons (127 neurons from 4 mice) across trials in response to five shock intensities. Cells were sorted based on their shock response to 0.90mA. **h**, Trial-averaged ΔF/F Ca_2+_ time-course from all recorded ITC_dm_ neurons. **i**, Mean shock responses. **j**, Proportions of significantly shock responsive neurons. **k**, Experimental scheme for reward consumption experiments. **l**, Example Ca_2+_ traces from the same neurons as in (**e**). **m**, Trial-averaged ΔF/F Ca_2+_ time-course aligned to reward consummation of all recorded ITC_dm_ from the representative mouse uses in (**f**). **n**,**o**, Trial-averaged ΔF/F Ca^2+^ time-course aligned to reward consummation of all recorded ITC_dm_ neurons. Response to sucrose and milk were averaged. In **n**, Cells were sorted based on their reward response. **p**, A scatter plot visualizing relationship between shock response and reward response.

To assess the properties of the same ITC_dm_ neurons in response to appetitive stimuli, after completion of the shock titration experiment, we food-restricted mice for five to seven days and performed another imaging session (**Fig. 1c**) where they were allowed to voluntarily consume two rewards, 20% sucrose and condensed milk, offered at random intervals (**Fig. 1k**; see Methods). Almost no ITC_dm_ neurons were excited by rewards, although some ITC_dm_ neurons showed inhibitory responses (i.e. reduced activity)(**Fig. 1l-n**), resulting in a slightly negative population average (**Fig. 1o**). There was no clear relationship between responses to appetitive and aversive stimuli (**Fig. 1p**), demonstrating that ITC_dm_ is a dedicated system for processing negative value.

We then performed a similar series of experiments on ITC_vm_ neurons (**Fig. 2a-c**). As previously shown in our data from fear conditioning experiments (Hagihara et al. 2021), most ITC_vm_ neurons were not excited by foot shocks at 0.65 mA (**Fig. 2d**). At the population level, a slightly inhibitory average shock response was observed (**Fig. 2g**). Interestingly, a small fraction (4.2%) of ITC_vm_ neurons showed shock offset-locked responses. Given most shock responsive ITC_dm_ neurons showed shock onset-locked responses (**Fig. 1f**), this offset response of ITC_vm_ could be a rebound excitation upon termination of inhibition by ITC_dm_ possibly associated with relief.

**Figure 2.**
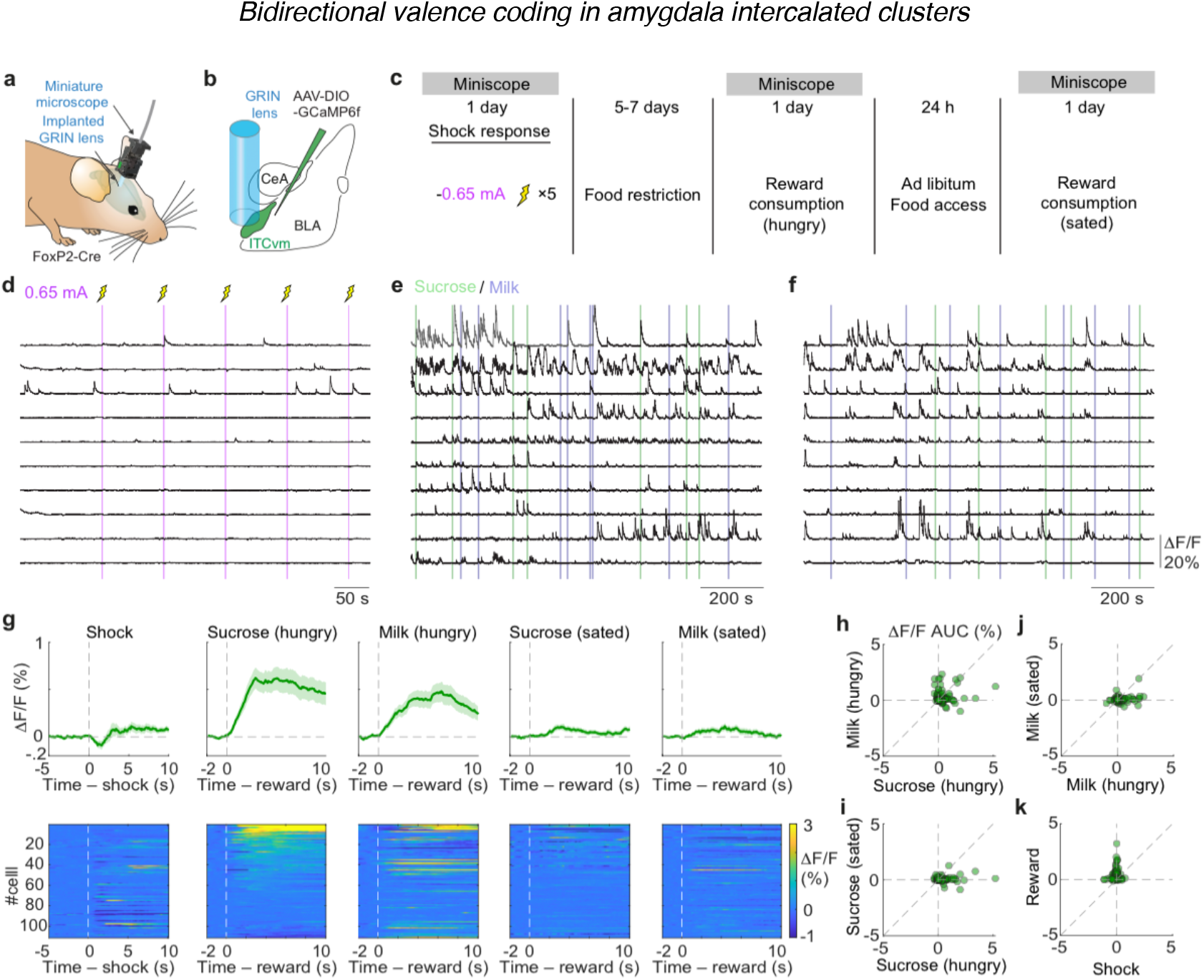
ITC_vm_ valence coding. **a**, Miniature microscope imaging in a freely-moving mouse. **b**, AAV encoding CAG-DIO-GCaMP6f was targeted to ITC_vm_ in FoxP2-Cre mice. GRIN lens implanted above injection site. **c**, Experimental time course. **d-f**, Example Ca_2+_ traces from 10 example neurons from shock (**d**), reward-hungry (**e**), reward-sated (**f**) sessions. **g**, Trial-averaged ΔF/F Ca^2+^ time-course aligned to shock and reward consummation of all recorded ITC_vm_ neurons (111 neurons from 5 mice). Cells were sorted based on their response to sucrose (second from left) and kept consistent for other heat plots. **h-k**, Scatter plots visualizing relationship between responses to sucrose-hungry and milk-hungry (**h**); sucrose-hungry and sucrose-sated (**i**); milk-hungry and milk-sated (**j**); shock and reward (average of sucrose-hungry and milk hungry) (**k**).

In stark contrast to ITC_dm_, which showed no excitatory reward responses, 42.3% of ITC_vm_ neurons showed significant reward responses in recording sessions after 5-7 days of food restriction (**Fig. 2e,g**). Some neurons showed selectivity between the two rewards (**Fig. 2h**). We adjusted the concentration of milk so that mice would show a similar degree of preference for the two rewards (Courtin et al. 2022) so that they would have similar positive values. Thus, this selectivity suggests that positive value is not the only factor that activates ITC_vm_ neurons. Instead, some ITC_vm_ neurons likely have a reward identity-specific positive value representation. To rule out the possibility that the observed ITC_vm_ reward responses reflect pure sensory qualities (taste) of the rewards rather than their positive value, we allowed the mice to have *ad libitum* access to food for 24 h after the first food-restricted recording session (**Fig. 2c**). Thus, in the second recording session, the mice were no longer hungry, and the value of the rewards would have been unselectively devalued. Indeed, although the mice still continued to consume rewards, the number of voluntary consummations was significantly reduced (9.6 ± 0.07 vs. 5.3 ± 1.1; *P* = 3.9 × 10^−4^, Wilcoxon signed-rank test, n = 5 mice). We found that neurons that responded to rewards in the first *hungry* session dramatically reduced their responses in the subsequent *sated* session (**Fig. 2f, g, j, i**), supporting the notion that the reward responses of ITC_vm_ neurons represent positive valence. There was no clear relationship between responses to appetitive and aversive stimuli (**Fig. 2k**), demonstrating that ITC_vm_ is a system dedicated to positive valence, in contrast to ITC_dm_, which is dedicated to negative valence (**Fig. 3a**).

**Figure 3.**
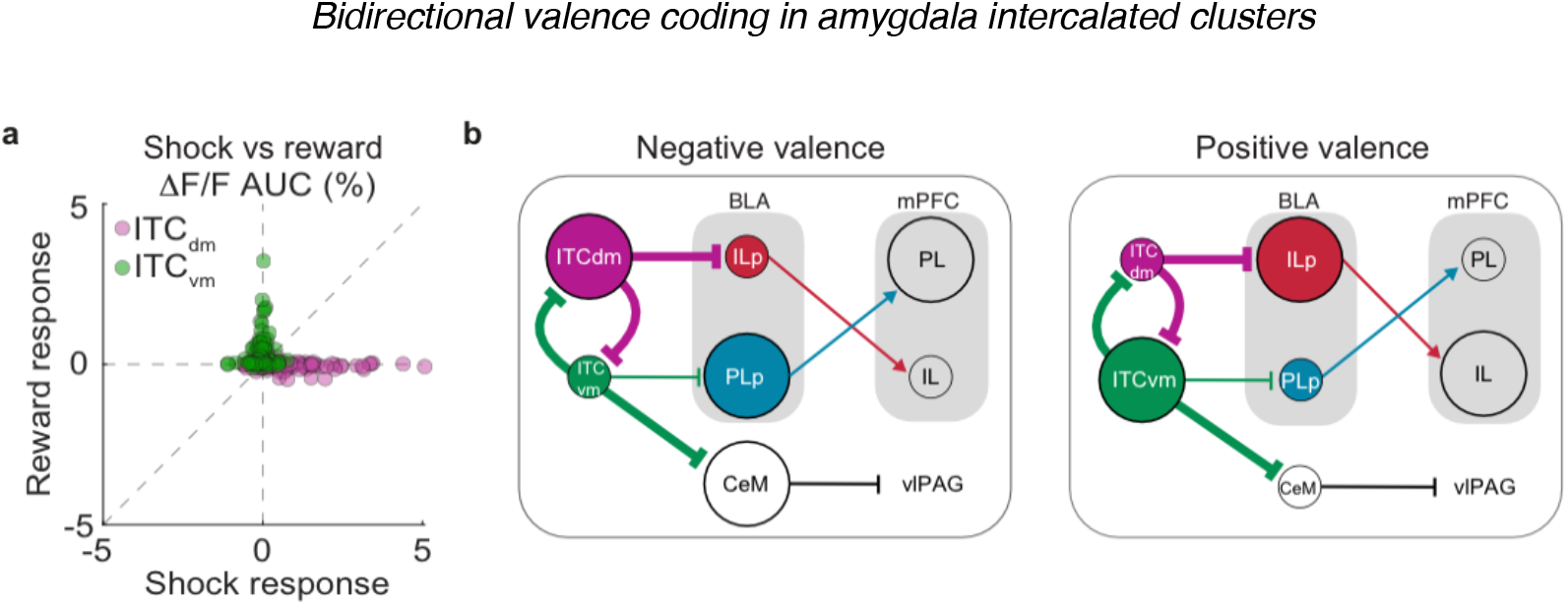
Summary. **a**, Shock and reward responses of ITC_dm_ and ITC_vm_. Data are duplicated from **Fig. 1p** and **Fig. 2k. b**, Summary scheme of the ITC function in orchestrating distributed circuits for positive and negative valence-specific behavioral selection. This incorporates previously identified functional connectivity between ITCs, BLA, medial prefrontal cortices (mPFC), medial central amygdala (CeM), and ventro-lateral peri-aquaductal gray (vlPAG) (Senn et al. 2014; Tovote et al. 2016; Hagihara et al. 2021; Massi et al. 2023).

## Discussion

In the current study, we found that the response properties of ITC_dm_ and ITC_vm_ are distinctly different – ITC_dm_ is active only upon presentation of negative valence stimuli, whereas ITC_vm_ responds only to stimuli with positive valence (**Fig. 3a**). Taken together with our previous work (Hagihara et al. 2021), the new findings demonstrate that the ITC system has a broader role beyond the acquisition and extinction of conditioned fear responses (**Fig. 3b**), and is a plausible neural circuit for implementing the opponent-process of motivation. To test this hypothesis, the following two questions should be addressed in the future: 1) whether ITC_dm_ shows a positive response to the omission of expected positive stimuli; 2) whether the activity of ITC_vm_ scales with the value of positive stimuli. Furthermore, Solomon and Corbit’s original theory included habituation and extinction upon repeated exposure to the affect-evoking US and CS (unconditioned and conditioned stimuli), respectively. Indeed, we found that repeated exposure to a CS in the absence of the US shifts the balance between ITC clusters and allows for fear extinction (Hagihara et al. 2021). It would be of great interest to test whether the same phenomenon can be observed in ITC_vm_ upon extinction of appetitively conditioned CSs.

In addition to ITCs, several distributed circuits have been identified that represent positive and/or negative valence, and are involved in generation of valence-specific behavior (Tye 2018). Some molecularly defined cell types show bidirectional representations: a conventional subpopulation of midbrain dopaminergic neurons respond bidirectionally to opposite valence – excited by positive valence and inhibited by negative valence (Schultz and Romo 1987; Ungless, Magill, and Bolam 2004; Brischoux et al. 2009; M. Matsumoto and Hikosaka 2009; Cohen et al. 2012; H. Matsumoto et al. 2016); corticotropin-releasing factor-releasing (CRF+) neurons in the paraventricular nucleus are inhibited upon exposure to stimuli with positive valence and inhibited by stimuli with negative valence (Jineun Kim et al. 2019). Some cell types, such as basal forebrain cholinergic cells, show unsigned responses to both positive and negative stimuli (Hangya et al. 2015). In addition, some brain regions such as ventral pallidum (Tachibana and Hikosaka 2012; Stephenson-Jones et al. 2020) and the basolateral amygdala (BLA) (Paton et al. 2006; Senn et al. 2014; Beyeler et al. 2016; O’Neill, Gore, and Salzman 2018; Kyriazi, Headley, and Paré 2020; Piantadosi et al. 2023) contain distinct neuronal populations that show opposite valence responses. The functional organization of the BLA is of particular interest because two spatially intermingled (but see (Joshua Kim et al. 2017)), but functionally opposite excitatory neuron populations, show mutual inhibitory dynamics through polysynaptic inhibition (Piantadosi et al. 2023) and/or external coordination mechanisms that orchestrate the two populations. In contrast to ITC neurons, which respond selectively to either positive or negative valence (**Fig. 3a**), a large fraction of BLA neurons respond to both (Beyeler et al. 2016; Piantadosi et al. 2023). Thus, it is tempting to hypothesize that mutual inhibition between the highly valence-selective ITC clusters carves the mutual inhibitory dynamics between the BLA excitatory neuron populations (**Fig. 3b**).

In both our previous (Hagihara et al. 2021) and current studies, we found that ITC_dm_ are functionally largely homogeneous, whereas ITC_vm_ neurons are more heterogeneous – about 20% of extinction neurons and 40% reward responding neurons. Their potential overlap is of interest and should be investigated in future experiments. Assessing the anatomical or molecular correspondence to the functional heterogeneity in ITC_vm_ would be a key step in understanding the functional organization of the ITC system. Although ITC neurons appear to constitute a “subclass” of GABAergic neurons (O’Leary et al. 2020), they could be further subdivided into several “types” (see (Zeng 2022) for definitions). Small populations such as ITCs are often overlooked in large-scale studies, and thus a targeted and tailored investigation would be necessary. Among the known interesting molecular features of ITCs is the expression of G protein-coupled receptors. In particular, Dopamine 1 receptors and µ-opioid receptors are highly enriched in ITCs and have been used as marker genes to identify ITCs (Fuxe et al. 2003; Jacobsen et al. 2006). How these neuromodulatory inputs, as well as glutamatergic and long-range GABAergic inputs contribute to ITC functions in concert with the inter-cluster monosynaptic inhibition awaits future investigation.

## Acknowledgements

We thank J. Y. Cohen and A. Holmes for reading and commenting on the manuscript, and discussion; J. Courtin for sharing the experimental set up for appetitive behavior; N. Karalis for discussion and inspiration; T. Eichlisberger for animal management; FMI animal facility, microscopy core facility (FAIM), and IT department for constant support. This work was supported by the European Research Council (ERC) under the European Union’s Horizon 2020 research and innovation program (grant agreement no. 669582), and a Swiss National Science Foundation core grant (310030B_170268) (to A.L.); by The Human Frontier Science Program (to K.M.H.); and by the Novartis Research Foundation.

## Author contributions

K.M.H. conceived the project, performed experiments, analyzed the data, and wrote the manuscript. A.L. supervised the project.

## Materials and Methods

### Mice

All animal procedures were performed in accordance with institutional guidelines and with current European Union guidelines and were approved by the Veterinary Department of the Canton of Basel-Stadt, Switzerland. FoxP2-IRES-Cre mice (JAX#030541) (Rousso et al. 2016) were used for Cre-dependent expression of viral vectors. For some experiments where a Cre-dependent expression system was not required, Arc-CreER mice (Guenthner et al. 2013) crossed with a tdTomato reporter line (Ai14) were used in addition to wild-type C57BL/6J mice. All the mice used in this study were reported in our previous publication (Hagihara et al. 2021). Mice were individually housed for at least two weeks before starting behavioural experiments. Animals were kept in a 12-h light/dark cycle with access to food and water *ad libitum* except for food restriction experiments. All behavioural experiments were conducted during the light cycle.

### Surgical procedures

All the procedures were performed as previously described in detail (Hagihara et al. 2021). Briefly, AAV2/5.CaMK2.GCaMP6f (for ITC_dm_) or AAV2/9.CAG.flex.GCaMP6f (for ITC_vm_) was unilaterally injected into the amygdala. For ITC_dm_: AP −1.4 mm (from bregma), ML −3.3 mm (from bregma), DV 4.4 mm (from pia); For ITC_vm_: AP −1.6 mm (from bregma), ML-3.1 mm (from bregma), DV 5.0 mm (from pia); After waiting at least 10 min for diffusion of the virus, a gradient-index microendoscope (ITC_dm_: φ1.0 x 9.0 mm, 1050-002179, Inscopix GRIN lens; ITC_vm_: φ0.6 x 7.3 mm, 1050-002177, Inscopix GRIN lens) was implanted.

### Deep brain calcium imaging and behavior experiments

Imaging experiments were performed as previously described in detail (Hagihara et al. 2021; Courtin et al. 2022). Briefly, two to six weeks after GRIN lens implantation, mice were habituated to the brief head-fixation on a running wheel for miniature microscope mounting for at least three days before the behavioural paradigm. Imaging data were acquired using nVista HD software (Inscopix) at a frame rate of 20 Hz. For individual mice, the same imaging parameters were kept across days.

Shock intensity titration experiments were performed in a clear square box with an electrical grid floor (Coulbourn Instruments) for foot shock delivery, placed in a light-coloured sound attenuating chamber with bright light conditions, and was scented and cleaned with 70% ethanol. A stimulus isolator (ISO-Flex, A.M.P.I.) was used for the delivery of direct current (DC) shock. Reward consumption experiments were performed in a box equipped with two lick-ports on the same wall as described previously (Courtin et al. 2022). Briefly, the lick-port was composed of an empty cylinder (made of POM) positioned horizontally with open window on the top where mice access liquids (open window: ellipse of 6 by 3 mm). Liquids were delivered in a receptacle inserted in the cylinder (receptacle: half-ellipsoid of 6 by 3 by 2 mm). The receptacle was made of aluminum to measure tongue contacts via an analog input board of Neural Recording Data Acquisition Processor system (OmniPlex, Plexon). Lick onsets were inferred off-line by detecting potential rise-times. Each lickport allowed delivery of either sucrose (20%) or sweetened condensed milk (15%, Régilait) solutions via TTL-controlled syringe pumps (PHM-107, Med Associates). Two rewards were provided ten times each with random intervals. Cameras (Stingray, Allied Vision) for tracking animal behaviour were also equipped in both chambers. Radiant Software (Plexon) was used to generate precise TTL pulses to control behavioural protocols and all the TTL signals including miniscope frame timings were recorded by Plex Control Software (Plexon) to synchronize behavioural protocols, behavioural tracking, and miniscope imaging. Upon completion of the behavioral experiments, mice were transcardially perfused with 4% PFA, and then, virus expression and GRIN lens implant sites were histologically verified (Hagihara et al. 2021).

### Statistical analyses and data presentation

All data are expressed as the mean ± standard error of the mean (SEM), unless stated otherwise. Two-sided Wilcoxon rank-sum test was used to determine significantly responding neurons. For paired comparison, we used Wilcoxon signed-rank test. For trend, Jonckheere-Terpstra test was used (**Fig. 1i**). Throughout the study, *P* < 0.05 was considered statistically significant. No statistical methods were used to pre-determine sample sizes, but our sample sizes are similar to those generally employed in the field.

### Data and code availability

Upon publication, the data that support the findings and custom-written codes used to analyse the data will be available at: https://data.fmi.ch/PublicationSupplementRepo/

## References

1. Hagihara, K. M. et al. Intercalated amygdala clusters orchestrate a switch in fear state. Nature (2021) doi:10.1038/s41586-021-03593-1.

2. Peterson, G. B., Ackilt, J. E., Frommer, G. P. & Hearst, E. S. Conditioned Approach and Contact Behavior toward Signals for Food or Brain-Stimulation Reinforcement. Science 177, 1009–1011 (1972).

3. Fanselow, M. S. & Bolles, R. C. Naloxone and shock-elicited freezing in the rat. J. Comp. Physiol. Psychol. 93, 736–744 (1979).

4. Handbook of anxiety and fear. Handbook of behavioral neuroscience. 517, (2008).

5. Blanchard, D. C., Griebel, G. & Blanchard, R. J. Mouse defensive behaviors: pharmacological and behavioral assays for anxiety and panic. Neurosci. Biobehav. Rev. 25, 205–218 (2001).

6. Tovote, P. et al. Midbrain circuits for defensive behaviour. Nature 534, 206–212 (2016).

7. Fadok, J. P. et al. A competitive inhibitory circuit for selection of active and passive fear responses. Nature 542, 96–100 (2017).

8. Pavlov, P. I. Conditioned reflexes: An investigation of the physiological activity of the cerebral cortex. Ann Neurosci 17, 136–141 (1927).

9. Tye, K. M. Neural Circuit Motifs in Valence Processing. Neuron 100, 436–452 (2018).

10. Gross, C. T. & Canteras, N. S. The many paths to fear. Nat. Rev. Neurosci. 13, 651–658 (2012).

11. Tovote, P., Fadok, J. P. & Lüthi, A. Neuronal circuits for fear and anxiety. Nat. Rev. Neurosci. 16, 317–331 (2015).

12. Herry, C. & Johansen, J. P. Encoding of fear learning and memory in distributed neuronal circuits. Nat. Neurosci. 17, 1644–1654 (2014).

13. Solomon, R. L. & Corbit, J. D. An opponent-process theory of motivation. I. Temporal dynamics of affect. Psychol. Rev. 81, 119–145 (1974).

14. Asede, D., Doddapaneni, D. & Bolton, M. M. Amygdala Intercalated Cells: Gate Keepers and Conveyors of Internal State to the Circuits of Emotion. J. Neurosci. 42, 9098–9109 (2022).

15. Rousso, D. L. et al. Two Pairs of ON and OFF Retinal Ganglion Cells Are Defined by Intersectional Patterns of Transcription Factor Expression. Cell Rep. 15, 1930–1944 (2016).

16. Guenthner, C. J., Miyamichi, K., Yang, H. H., Heller, H. C. & Luo, L. Permanent genetic access to transiently active neurons via TRAP: targeted recombination in active populations. Neuron 78, 773–784 (2013).

17. Courtin, J. et al. A neuronal mechanism for motivational control of behavior. Science 375, eabg7277 (2022).

18. Ghosh, K. K. et al. Miniaturized integration of a fluorescence microscope. Nat. Methods 8, 871–878 (2011).

19. Schultz, W. & Romo, R. Responses of nigrostriatal dopamine neurons to high-intensity somatosensory stimulation in the anesthetized monkey. J. Neurophysiol. 57, 201–217 (1987).

20. Ungless, M. A., Magill, P. J. & Bolam, J. P. Uniform Inhibition of Dopamine Neurons in the Ventral Tegmental Area by Aversive Stimuli. Science 303, 2040–2042 (2004).

21. Brischoux, F., Chakraborty, S., Brierley, D. I. & Ungless, M. A. Phasic excitation of dopamine neurons in ventral VTA by noxious stimuli. Proceedings of the National Academy of Sciences 106, 4894–4899 (2009).

22. Matsumoto, M. & Hikosaka, O. Two types of dopamine neuron distinctly convey positive and negative motivational signals. Nature 459, 837–841 (2009).

23. Cohen, J. Y., Haesler, S., Vong, L., Lowell, B. B. & Uchida, N. Neurontype-specific signals for reward and punishment in the ventral tegmental area. Nature 482, 85–88 (2012).

24. Matsumoto, H., Tian, J., Uchida, N. & Watabe-Uchida, M. Midbrain dopamine neurons signal aversion in a reward-context-dependent manner. Elife 5, (2016).

25. Kim, J. et al. Rapid, biphasic CRF neuronal responses encode positive and negative valence. Nat. Neurosci. 22, 576–585 (2019).

26. Hangya, B., Ranade, S. P., Lorenc, M. & Kepecs, A. Central Cholinergic Neurons Are Rapidly Recruited by Reinforcement Feedback. Cell 162, 1155–1168 (2015).

27. Tachibana, Y. & Hikosaka, O. The primate ventral pallidum encodes expected reward value and regulates motor action. Neuron 76, 826–837 (2012).

28. Stephenson-Jones, M. et al. Opposing Contributions of GABAergic and Glutamatergic Ventral Pallidal Neurons to Motivational Behaviors. Neuron 105, 921-933.e5 (2020).

29. Paton, J. J., Belova, M. A., Morrison, S. E. & Salzman, C. D. The primate amygdala represents the positive and negative value of visual stimuli during learning. Nature 439, 865–870 (2006).

30. Senn, V. et al. Long-range connectivity defines behavioral specificity of amygdala neurons. Neuron 81, 428–437 (2014).

31. Beyeler, A. et al. Divergent Routing of Positive and Negative Information from the Amygdala during Memory Retrieval. Neuron 90, 348– 361 (2016).

32. O’Neill, P.-K., Gore, F. & Salzman, C. D. Basolateral amygdala circuitry in positive and negative valence. Curr. Opin. Neurobiol. 49, 175– 183 (2018).

33. Kyriazi, P., Headley, D. B. & Paré, D. Different Multidimensional Representations across the Amygdalo-Prefrontal Network during an Approach-Avoidance Task. Neuron 107, 717-730.e5 (2020).

34. Piantadosi, S. C. et al. Holographic stimulation of opposing amygdala ensembles bidirectionally modulates valence-specific behavior via mutual inhibition. Neuron (2023) doi:10.1016/j.neuron.2023.11.007.

35. Kim, J., Zhang, X., Muralidhar, S., LeBlanc, S. A. & Tonegawa, S. Basolateral to Central Amygdala Neural Circuits for Appetitive Behaviors. Neuron 93, 1464-1479.e5 (2017).

36. O’Leary, T. P. et al. Extensive and spatially variable within-cell-type heterogeneity across the basolateral amygdala. Elife 9, e59003 (2020).

37. Zeng, H. What is a cell type and how to define it? Cell 185, 2739–2755 (2022).

38. Fuxe, K. et al. The dopamine D1 receptor-rich main and paracapsular intercalated nerve cell groups of the rat amygdala: relationship to the dopamine innervation. Neuroscience 119, 733–746 (2003).

39. Jacobsen, K. X., Höistad, M., Staines, W. A. & Fuxe, K. The distribution of dopamine D1 receptor and mu-opioid receptor 1 receptor immunoreactivities in the amygdala and interstitial nucleus of the posterior limb of the anterior commissure: relationships to tyrosine hydroxylase and opioid peptide terminal systems. Neuroscience 141, 2007–2018 (2006).

40. Massi, L. et al. Disynaptic specificity of serial information flow for conditioned fear. Science Advances 9, eabq1637 (2023).

